# Acoustic markers of negative arousal in lambs: evidence from behavioural and eye thermal profiles

**DOI:** 10.1101/2025.07.29.667358

**Authors:** Stefania Celozzi, Zoe Miot, Paul Renaud-Goud, Silvana Mattiello, Elodie F. Briefer, Avelyne S. Villain

## Abstract

Vocalizations are known indicators of emotional arousal in animals, but validation using simultaneously collected physiological and behavioural measures remains limited to a few species. This study investigated sheep vocal expression of negative emotional arousal using stress-related behaviours and eye temperature as non-invasive arousal indicators. To this aim, twenty lambs underwent a short-term isolation test with two phases aimed at eliciting different levels of arousal in response to separation from conspecifics: partial isolation, where lambs maintained visual, acoustic and tactile contact with conspecifics through a fence, and full isolation (complete separation). During full isolation, lambs expressed higher bodily activation—spending more time running, jumping, and changing state behaviours—and produced more open-mouthed bleats (321 *vs* 27) than in partial isolation, validating higher arousal. Eye temperature also increased from partial to full isolation (however only in small lambs and not large ones). Calls emitted in full isolation were characterised by higher frequencies, were less tonal (more chaos) and had shorter durations. When combining behavioural and physiological assessment of arousal and testing their impact on the spectro-temporal structure of vocalizations, we found that bodily activation, but not eye peak temperature, impacted on the frequency distribution and tonality of the calls. Call duration increased with eye temperature, but only in lambs expressing high bodily activation, while the mean of the second formant increased with eye temperature in smaller but not larger lambs. Overall, lamb vocalizations indicate arousal and are correlated with bodily activation, while co-variations with physiological measures depended on behaviour and individual traits.

**Highlights:**

- Animal calls indicate emotional arousal, but few species have validated measures
- Lamb calls became higher in frequency, less tonal and shorter with negative arousal
- Calls indicated emotional arousal and were correlated with bodily activation
- Changes in calls with eye temperature depended on behaviour and individual traits

## 1. Introduction

Emotions are short-term multi-component responses to rewarding or punishing stimuli or events (Mendl & Paul, 2020). Among the theories developed to quantify emotions in non-human animals (hereafter ‘animals’), the dimensional theory is one of the most widely applied. According to this theory, emotions are characterised by their valence, which can be positive (pleasant) or negative (unpleasant), and arousal (i.e., bodily activation) (Russell, 1980; Mendl et al., 2010; Bradley et al., 2001). These two dimensions can be assessed using behavioural, physiological and cognitive indicators, such as postural changes (Reefmann et al., 2009a; de Oliveira & Keeling, 2018; Briefer et al., 2015a; Rius et al., 2018), cardiorespiratory measures (Reefmann et al., 2009b; Davies et al., 2014), temperature variations (Proctor & Carder, 2015), or judgement bias tasks (Baciadonna et al., 2016).

Infrared thermography is a technique that indirectly assesses body temperature through emitted infrared radiation. In recent years, this technique has been increasingly explored as a non-invasive tool to assess animal emotions (Travain & Valsecchi, 2021). Temperature changes appear particularly useful for assessing arousal but they can vary across body regions. For example, increased arousal is associated with decreased nasal temperature in marmosets (Ermatinger et al., 2019), decreased tail and paw temperature in rats (Vianna & Carrive, 2005), and increased eye temperature in dogs (Rigterink et al., 2018). These findings indicate that thermographic responses are region- and species-specific, requiring careful interpretation.

Alongside physiological responses, vocalizations are also valuable non-invasive indicators of animal emotions across taxa (Villain & Briefer, in production; Briefer, 2012; Briefer, 2020). Many vocal features change consistently with increasing arousal across species, whereas vocal correlates of valence are often species specific (Briefer, 2020). In high-arousal contexts, calls are typically louder, with higher frequencies and greater variability in fundamental frequency (hereafter ‘*fo*’). Because of these cross-species regularities, vocalisations are considered promising universal indicators of arousal (Briefer, 2020).

Sheep (*Ovis aries*) are gregarious prey animals, and separation from conspecifics is therefore stressful and fear-inducing (Cockram, 1994). Behavioural responses to social stress include changes in body and ear posture, mobility and rate of vocal activity (Ligout et al., 2011; Engeldal et al., 2013; Boissy et al., 2011; Han et al., 2024). Thermography has been shown to be useful for assessing sheep negative emotions; transport-related stress increases core temperature, while social isolation reduces ear pinna temperature due to vasoconstriction (Ingram et al., 2002; Lowe et al., 2005). Several studies additionally report increased eye temperature in response to stressors such as tethering, shearing, and fear-inducing stimuli (Cannas et al., 2018; Arfuso et al., 2022; Comin et al., 2024; Ingram et al., 2002; Lowe et al., 2005).

Sheep vocalisations serve several functions, including maternal bonding (Sèbe et al., 2007), mutual recognition (Papadaki et al., 2021), and group cohesion (contact calls), and can signal stress (Dwyer, 2008). Sheep produce low-pitched bleats (mouth closed), high-pitched bleats (mouth open) and mixed bleats (starting with a closed mouth and finishing with opened mouth). Low-pitched bleats or “rumble” sounds are typically produced by mothers near their lambs or by rams during courtship (Dwyer, 2008). High-pitched bleats are produced in high-arousal situations (Papadaki et al., 2021), which can be both positive (e.g., receiving a reward), and negative, (e.g., isolation and separation (Dwyer, 2008). However, little is known about how the spectro-temporal structure of sheep vocalizations varies with emotional arousal. Existing studies report increased *fo* contour (Sèbe et al., 2012) and a shift toward lower spectral energy (i.e., lower frequencies; Sèbe et al., 2012; Engeldal et al., 2013), with increasing negative arousal. Papadaki et al. (2021) also found higher *fo*-related features (minimum and mean *fo* and *fo* disturbance), and higher frequency distributions in ewes without lambs (dry period) compared to those with lambs (suckling period), possibly reflecting higher stress or hormonal differences. With the exception of Sèbe et al. (2012), other studies did not simultaneously measure independent indicators of arousal, leaving the vocal correlates of this dimension insufficiently understood.

This study aimed to address these gaps by examining vocalizations as indicators of negative emotional arousal in sheep, using behavioural and physiological measures as independent indicators of arousal. Lambs were subjected to a two-phase isolation test (partial and full), designed to induce a graded increase in negative arousal. Based on the literature, we predicted increased bodily activation and vocal rate, along with longer bleats of higher frequencies, and increased eye temperature when sheep went from partial to full isolation. Additionally, we expected vocal parameters to correlate with behavioural and thermal proxies of arousal, regardless of the lamb biometry.

## 2. Materials and methods

### 2.1. Animals and housing conditions

The study was carried out on a commercial meat sheep farm in North Zealand, Denmark, from April to May 2023. The flock was composed by 110 sheep (*Ovis aries*) of different breeds (Frisian milksheep, Lacaune, Charolais, Zwartbles, Texel, Spelsau Lleyn Suffolk and Dorset). Among these sheep, 58 were lambs aged between four and five months at the start of the experiment, and 52 were ewes up to eight years of age. The lambs were still unweaned and remained with their mothers throughout the study. All the animals were housed in the same barn of 200 m^2^ on a deep-litter straw bedding and had daily access to pasture from 8 am to 4 pm, accompanied by the farmer and sheepdogs. The pasture was composed of two areas of 15,100 m^2^ and 36,200 m^2^. All animals were managed in the same way in terms of feeding, which was calibrated to the nutritional requirements of adult sheep at approximately 320 g of barley per animal, divided in two daily administrations at 8.30 am and 3 pm. All sheep had *ad libitum* access to water.

The animals were naive to any experiment and were used for two subsequent experiments. In the other experiment (not described here), sheep were exposed to a playback experiment in the field, involving no previous manipulation. This playback experiment occurred after the tests presented in this study.

### 2.2. Experimental procedure

#### 2.2.1 Individual identification and habituation

All sheep were marked using specific sheep spray (Agrihealth Quickdry Sheep Marker Spray) according to a unique colour code applied to the back and flanks of each animal. For the need of the other experiment running at the same time, the flock was divided into two homogeneous groups in terms of numbers, keeping the lambs together with their mothers. The barn was divided in two equal parts, and each group was assigned to a pasture field. A habituation phase of two days was adopted to let the animals familiarize themselves with the division of the flock, the location of pasture field they were assigned to, and the position of the feeders in the fields. The sheep were also habituated to the presence of two experimenters, who walked for 10 min twice a day in the fields for two days, maintaining approximately a distance of thirty meters from the animals. In addition, the sheep were habituated to the equipment: a hide (supplementary material, figure S1), which was installed in the barn to allow the experimenters to observe the sheep without visual disturbance, and a metal pen.

#### 2.2.2 Isolation tests

The isolation tests started after the habituation phase. A total of twenty lambs were randomly selected from the flock for the isolation tests. Eligibility criteria required that individuals were juveniles, healthy, and naïve to previous experimentation. No selection was made based on sex, breed, body size, or behavioural tendencies; thus, the final sample (11 females, 7 males and 2 of unknow sex) reflected the natural variation within the lamb population. This procedure aligns with the STRANGE framework, ensuring that the study subjects were representative of the genetic experiential and social diversity within the target population (Webster & Rutz, 2020).

The tested lambs were isolated from conspecifics in the morning (08:21 (hh:min) ± 00:12) before the flock was moved to the pasture. Each day, a randomly selected lamb was gently restrained by two experimenters and placed in a metal pen within the barn where the animals were normally housed. The pen was shaped like an irregular polygon, approximately 1.43 m high and with a surface area of 2 m^2^ (supplementary material, figure S1). The isolation test was divided into two phases: 1) the lamb had visual, vocal and physical access to its group members through the grid of the pen (partial isolation); 2) the group was guided toward the pasture field by one experimenter, while the tested lamb remained in the isolation pen in the absence of any physical, visual or vocal contact with group members (full isolation). Each phase lasted 4 minutes, for a total duration of 8 minutes. At the end of the test, the subject was allowed to reunite with the flock at pasture. Each lamb was tested only once, with only one animal tested per day, for a total of twenty experimental days.

### 2.3. Behaviour recording and scoring

Lambs’ behaviour during the tests was video recorded using a GoPro Hero9 Black (GoPro Inc, CA, USA). Videos were later scored by a trained observer, who continuously recorded lambs’ behaviour during the partial and full isolation tests (total observation time of 8 min per lamb), using an ethogram created with BORIS software (Friard & Gamba, 2016), which included both state and point events (Table 1). Ten percent of the videos were scored twice to assess intra-observer reliability, which was quantified using Cohen’s Kappa and ranged from 0.731 to 0.836.

**Table 1.** Ethogram of the behaviours and postures scored during the isolation tests and their description.

| Scored behaviour | Behaviour type | Description |
| --- | --- | --- |
| **Bodily activation** |  | **Static (standing + lying down) and dynamic (walking + running) postures** |
| Standing | State | The lamb stands on its four legs and remains still for at least two seconds |
| Walking | State | The lamb slowly moves forward or backward |
| Running | State | The lamb moves forward with a fast gait, lifting one limb before the other is supported |
| Head movements | Point | All changes (i.e., fast and slow) in the position of the lamb's head in any direction |
| Bar head touches | Point | The lamb sticks its head between the bars of the pen or hits them |
| Jump/rearing on the hind legs | Point | The lamb is off the ground with all legs raised/the lamb is standing on its hind legs |
| Lying down | State | The lamb is in sternal or lateral decubitus (abdomen in contact with straw bedding) |
| **Ears position<sup>1</sup>** |  | **The position of the lamb's ears in relation to the orientation of the auricles** |
| Open | State | The inner and outer sides are not visible, the inner side is parallel to the floor, the observer can see the full ears |
| Flat | State | The inner sides are visible by an observer placed in front of the animal |
| Hidden | State | The inner sides are not visible, the outer sides are visible at the root of the ear, the ears cannot be fully seen by the observer |
| Asymmetric | State | One inner side visible, one not |
| Unidentifiable | State | The position of the lamb's ears cannot be classified because at least one of them is not visible |
| **Tail position** |  | **The position of the lamb's tail** |
| Upward | State | The lamb raises its tail up above a 45-degree angle |
| Down | State | The lamb has its tail down, below a 45-degree angle |
| Arched upward | State | The lamb keeps its tail arched over its rump |
| Shaking | State | The lamb moves its tail rapidly in any direction |
| At 45 degrees | State | The lamb has its tail raised at a 45-degree angle |
| Unidentifiable | State | The position of the lambs' tail is not visible |
| **Vocalisation** |  | **The lamb is bleating** |
| Open mouth | Point | The lamb bleats with its mouth open or begins to bleat with its mouth closed and ends with its mouth open |
| Closed mouth | Point | The lamb bleats with its mouth closed |
| **Excreting behaviours** |  | **The lamb is urinating or defecating** |
| Urinate | Point | The lamb is urinating |
| Defecate | Point | The lamb is defecating |
| **Social behaviours** |  |  |
| Social licking | Point | The lamb is licking one or more members of the audience |
| Social sniffing | Point | The lamb is sniffing one or more members of the audience |
| Presence of the audience | State | The lamb is considered in proximity to the audience when at least one group member is within one sheep's length away |
| Absence of the audience | State | The lamb is considered not in proximity to the audience when no group members are within one sheep's length away |
| Sheep to sheep contact | State | The lamb is in close physical contact with at least one audience member |
| Number of subjects in contact | Specifier | The number of subjects with whom the lamb is in close physical contact |

The percentage of time spent performing state behaviours was calculated as the duration of the behaviour divided by the total time during which the behaviour was visible, multiplied by 100. The rate of occurrence of point events was calculated as the number of occurrences divided by the time during which the behaviour was visible (expressed as events per minute). Biometric differences can affect both the acoustic parameters (Taylor & Reby, 2010) and the thermoregulatory process (Kleiber, 1947). Unfortunately, due to technical reasons, the weight of the animals, could not be measured as a proxy of body size. Hence, we estimated lambs’ height by visualizing the video recordings in the standard test environment and using the height of a pen bar as a reference measure. Lambs’ height was used as a proxy of lambs’ size, which was classed as “small” (height < 52 cm) or “large” lambs (height > 52 cm).

### 2.4 Thermal profiles

Thermal images of the lambs in partial and full isolation were taken with a Hikmicro G60 handled thermal camera by using the following parameters: emissivity = 0.98, reflected temperature = 25°C and distance = 4 m. The reflected temperature was kept constant at 25 °C, because our analysis relied on relative changes in eye temperature between partial and full isolation; the short 8-minute window (4 minutes during partial isolation and 4 minutes during full isolation) minimised the influence of ambient temperature and humidity, making daily adjustments unnecessary. To avoid influencing the subject’s emotional response, the experimenter took the images from the hide, through a small opening. Thermal images were collected *ad libitum* during both phases of the test (partial and full isolation, n = 3010 images in total), they were manually inspected to determine their quality, resulting in the selection of 707 high-quality pictures (435 were acquired during partial isolation and 272 during full isolation). Eye areas from these pictures (one or two eyes per picture, depending on the lamb position, and consequently on eye visibility) were manually defined in the Hikmicro software. Outliers, located outside the mean +-3sd range, were removed, as they likely derived from false calculations of eye temperature, resulting in 755 eye areas retained for subsequent analyses. The maximum temperature of each eye was used for statistical analysis to avoid the impact of how the area boundaries were defined on the computation of the mean temperature (Mason, 2023). Finally, a delta time value was calculated between the time when the thermal images were taken and the start of each phase of the test, in order to have an information about the effect of the time on thermal eye profiles.

### 2.5 Acoustic recordings and analyses

Vocalizations produced by the lambs during the isolation tests were recorded with an omnidirectional microphone (Sennheiser MKH20) placed at approximately 3.4 m in front of and 2.4 m above from the isolation pen, mounted on a metal rod fixed to the barn structure. The microphone was connected to a recorder (Marantz PMD661, sampling frequency: 44100 Hz and 16-bit WAV file format). Vocalizations were manually labelled according to the call type (low or high bleats) and checked for sound quality: only clean calls, *i.e.* having no overlapping sounds and a correct Signal to Noise Ratio (SNR), were retained for analysis (‘get_SNR_dir’ function, *SoundChunk* R-package; Villain & Renaud-Goud, 2023). The inspection of the Signal to Noise Ratio (SNR) revealed that calls (n = 464) were louder by 10.1 ± 0.56 dB compared to background noise. Only High Bleats (HB) were included in the statistical analysis, as the incidence of low bleats was very low (1.56% of total recorded vocalizations). After call selection, acoustic parameters were extracted on the remaining clean HB (n = 348; 17.4 ± 9.8 calls per individual) (‘extract_chunk_features_dir’ function, *Soundchunk* R, *tuneR* and *Seewave* R packages; Ligges et al., 2023; Sueur et al., 2008; see supplementary Table S1). The following automatic computations were applied, after inspection of the frequency range of interest, frequency range of the fundamental frequency (*fo*) and number of putative formants, on averaged mean spectra (supplementary figure S2). The sound was bandpass-filtered between 150 and 6000 Hz (“hanning” window type, FFT window length = 1024 pts and overlap = 50%). The duration of the calls was determined using a 3% amplitude threshold on the smoothed envelop (window length = 4000 pts, overlap = 90%). The extracted acoustic features related to the mean spectrum were the Median, Inter Quartile Range (IQR) and the Wiener entropy (sfm). The Dominant Frequency (supplementary figure S3A) was computed on each sliding temporal window (FFT window length = 1024 pts and overlap = 50%) and detected with a 10% amplitude threshold, by searching for the loudest frequencies in each window. The average (Dominant frequency mean) and the standard deviation (Dominant frequency sd) were extracted as descriptors of the dominant frequency and its variability. The fundamental frequency (*fo*) was detected by filtering the calls with a frequency range of 150-350 Hz and by searching for the loudest frequencies within this band (supplementary figure S3A). The average (*fo* mean) and the standard deviation (*fo* sd) were extracted as descriptors of the fundamental frequency and its variability. Finally, formant analysis was performed by searching for three main formants between 350 and 6000 kHz (supplementary figure S3B). Based on the detection of three peaks on a mean spectrum computed with FFT window length of 256 pts, the relative amplitude of formants F1 and F2 were extracted. Three frequency intervals were then computed to extract the frequency in each frequency interval (Relative Amp. F2, F3), corresponding to the three putative formants. For each frequency interval, the calls were filtered and the average of dominant frequencies computed over the duration of the call was extracted (mean F1, F2, F3). Each acoustic parameter and its definition and unit are available in supplementary table 1.

### 2.6 Statistical analysis

All statistical analyses were all performed in R 4.4.2. We first validated each indicator (behavioural, thermal, acoustic) by examining its variation from one phase of the test to the other within individuals, and subsequently explored their co-variation by integrating all indicators into a within-subject repeated-measures design.

#### 2.6.1 Behavioural response

After inspection of zero inflated data on rates and percentages of time spent expressing the scored behaviour, the ear and tail positions and excreting behaviours were excluded from further behavioural analysis, due to their non-symmetrical distribution despite the application of multiple transformation techniques. Three behavioural variables were calculated from the catalogued behaviours; 1) state behaviours changes, namely the number of transitions between standing, walking, running and lying down behaviours per time unit, 2) the percentage of time being in static posture (aggregating standing and lying down behaviours), and 3) a Social Score during the partial isolation phase, namely the sum of the following measures for each lamb: the presence or absence of the audience (scored 1 and 0, respectively), the number of social sniffing and licking, the number of body contacts with at least one audience member, and the total number of individuals in contact. The social score was initially included in the models, but was subsequently removed as it had no effect on the response variables. The behavioural variables used for analysing the behavioural response were therefore: the rate of jumps, state behaviours changes, bar head touches, head movements, vocalisations, quantified as the number of calls emitted per minute (hereafter referred to as *vocal dynamic*), and percentage of time spent walking, running or being static (Dray & Dufour, 2007). After inspection of symmetrical distributions of behavioural variables and transformations when needed (see Table 2), a Principal Component Analysis (PCA) was performed on the abovementioned behavioural variables to generate composite behavioural scores and reduce the number of correlated response variables (‘dudi.pca’ function, ‘ade4’ R package). All principal components (PCs) with an eigenvalue greater than one were retained for further analyses and used as behavioural composite scores (behPCs).

**Table 2.** Results of the Principal Component Analysis performed on the behavioural dataset, showing the parameters loadings on PC1 and PC2 (relative contribution for each behaviour to the PCs). The high parameters’ loadings (i.e., > 50 or < −50) are in bold.

| Behavioural parameters | PC1 | PC2 |
| --- | --- | --- |
| Jump rate<sup>1</sup> | **70.7** | 0.01 |
| Percent of total time in static postures<sup>2</sup> | **-65.5** | -0.01 |
| Percent of total time walking<sup>2</sup> | 5.08 | **-88.3** |
| Percent of total time running | **76.7** | 11.5 |
| Rate of state behaviours changes | **74.3** | -2.17 |
| Vocal dynamic<sup>1</sup> | **80.1** | 4.91 |
| Rate of bar head touches | **52.0** | 0.58 |
| Rate of head movements | 43.2 | -12.0 |

1 Square root-transformed variable
2 Ln-transformed variable

To explain variation in the lamb’s behavioural scores (behPC1, behPC2), which were entered as response variables in a Linear Mixed-Effects Model (LMM: ‘lmer’ function,‘lme4’ R package; Bates et al., 2015), we entered the following fixed effects in our model: the ‘phase of the test’ (two levels: partial *vs* full isolation), the size of the lamb (two levels: ‘small’ *vs* ‘large’) and the interaction between these two factors. The identity of the lamb was used as random factor to take into account the within subject design.

*Model 1: behPC1/behPC2 ∼ Phase of the test*Lamb size + (1|Subject)*

#### 2.6.2 Thermal response

Considering natural within-subject physiological variations in eye temperature, the effect of full isolation on eye maximum temperature was assessed by computing a coefficient of reaction (CR) during the full isolation compared to the partial isolation of the same individual, used as the ‘baseline’. The CR was computed as follows:

*temperatureCR=(T - baseline)/baseline*

*where:*

*T = maximum eye temperature in sample*

*baseline = average of maximum eye temperature in partial isolation*

Using this computation, positive or negative *temperatureCR* during the full isolation phase would be the result of, respectively, increasing or decreasing temperature compared to individual baseline values during partial isolation. The larger the absolute value of the *temperatureCR* was, the stronger the variation from partial to full isolation. Model 2 was built to test the effect of the phase of the test on *temperatureCR*, while using the size of the lamb as interacting fixed effect. In addition, the centred and reduced delta time to the start of the phase (Zdelta) was also added as interactive covariate to the phase of the test, in order to control for the effect of time within each phase on thermal eye profile. Lastly, the identity of the subject was included in the model as a random effect to control for repeated measurements.

*Model 2: temperatureCR ∼ Phase of the test* (lamb size + Zdelta) +(1|Subject) data = all eye max temperature data points*

#### 2.6.3 Acoustic response

Similarly to the behavioural variables, composite acoustic scores were computed to describe the general structure of the vocalisations while decreasing the number of correlated variables. A PCA was performed on the set of acoustic features listed in Table S1 (‘dudi.pca’ function, ‘ade4’ R package, Dray & Dufour, 2007), after inspecting their symmetrical distribution. All principal components with an eigenvalue greater than one were retained to be used as descriptors of the spectrotemporal structure of lamb bleats (acPCs). Furthermore, to align the direction of the acoustic feature loadings with the conventional representation of arousal in the dimensional theory of emotions (Mendl et al., 2010)—i.e., increasing from bottom to top—the signs of the first two principal components were inverted.

The comparison between the calls produced in partial and in full isolation was investigated with a LMM (‘lmer’, ‘lme4’ R package; Bates et al., 2015). The following fixed factors were included the model: the phase of the test (partial or full isolation) and the size of the lamb (small or large), as well as their interaction (Model 3). The identity of the subject was included in the model as a random effect to control for repeated measurements. Since not all lambs produced calls during the partial isolation, this analysis was performed twice, first considering all the lambs and then considering only the lambs that had produced calls in both phases (6 animals: N = 27 calls during partial isolation, and N = 128 calls during full isolation). Since the models using the entire dataset and the subset yielded similar results, only the model based on the entire dataset is presented in the results and used for further analysis (see Table S11 in supplementary material for statistical results referred to the subset).

*Model 3: acPCs ∼ phase of the test*lamb size + (1|Subject), data = all calls*

#### 2.6.4 Impact of behavioural and physiological indicators on vocalisations

To estimate co-variation between the acoustic structure of high bleats (acPCs), the behavioural response (behPC1) and the thermal profiles (eye maximum temperature) in response to isolation, a full model was built using each composite acoustic scores (acPCs) as response variables (Model 4), and the first behavioural score (behPC1), describing the bodily activation of the lamb in response to full isolation (see model 1) as a proxy of arousal, as a continuous fixed effect. The size of the lamb, which affected both the acoustic scores and the eye thermal profile, was used as factorial fixed effect (see models 2 and 3). Considering the increase in eye maximum temperature in response to full isolation (see model 3) and the effect of time during the phase on temperature variation (see model 3), the peak eye temperature per lamb and per phase was used as a further continuous fixed effect in the model. Since the number of temperature points per phase and lamb ranged from one to 46, and since 90% of the data was composed of four or more temperature points per phase and subject, we decided to average the four highest temperatures per lamb and per phase to obtain a more representative value of the eye peak temperature than using only one point per subject and phase. The interaction between behPC1 and eye peak temperature, as well as the interaction between lamb size and eye peak temperature, were also included to test for co-variation. Lastly, the individual identity of the subject was included in the model as a random effect to control for repeated call measurements.

*Model 4: acPCs*∼ *peak eye temperature * (behPC1 + lamb size) + (1|Subject), data = all calls*.

#### 2.6.5 Model fit and extraction of p - values (for all models)

After inspection of the distribution of the residuals of the models, a type-II analysis-of-variance was performed on each model to extract the corresponding p values (‘Anova’ function, ‘car’ R package, Fox & Weisberg, 2018). When significant interactions between fixed effects were found, pairwise comparisons were computed using ‘emmeans’ and ‘emtrends’ function from ‘emmeans’ R package (Lenth, 2016), with Tukey correction for factors and ‘emtrends’ function for interacting covariates. When significant interactions were found in model 4, comparisons were carried out by fixing some values of the covariate behPC1 (mean, mean - sd, mean + sd).

### 3. Ethical statement

This study was approved by the Animal Ethics Institutional Review Board at the Faculty of Health and Medical Sciences of the University of Copenhagen (official number: 023-03-BehaVEco-007A). As social isolation induces a negative affective state in the lambs, both isolation phases lasted only 4 minutes each. The isolation took place in a familiar environment, and any negative effect of this procedure on the sheep was not expected to last beyond the end of the test. Five isolation trials were interrupted because the lamb being tested escaped.

## 4. Results

### 4.1 Behavioural response across partial and full isolation

The Principal Component Analysis (PCA) performed on the behavioural variables resulted in two principal components with eigenvalues > 1. The first principal component (behPC1, 58.4% of explained variance) was strongly positively correlated with vocal dynamic, time spent running, rate of state behaviours changes, jumps and bar head touches, and strongly negatively correlated with time spent in static postures (standing + lying). Overall, behPC1 thus reflected bodily activation. The second principal component (behPC2, 14.9% of explained variance) was strongly negatively correlated with the percentage of time spent walking.

BehPC1 scores significantly increased, indicating an increase in bodily activation, from partial to full isolation (Chisq = 96.3; df = 1, p<0.001; figure 1) (Table 2). No effect of the size of the lamb on behPC1 scores was observed (Chisq = 0.13; df = 1, p = 0.71; supplementary Table2. Similarly, no effects of the phase of the test nor of the size of the lamb were found on behPC2 (Chisq = 2.34; df = 1, p = 0.13; Chisq = 2.92; df = 1, p = 0.09, respectively; supplementary Table 2. Only behPC1 was therefore used as proxy of bodily activation for further analysis.

**Figure 1.**
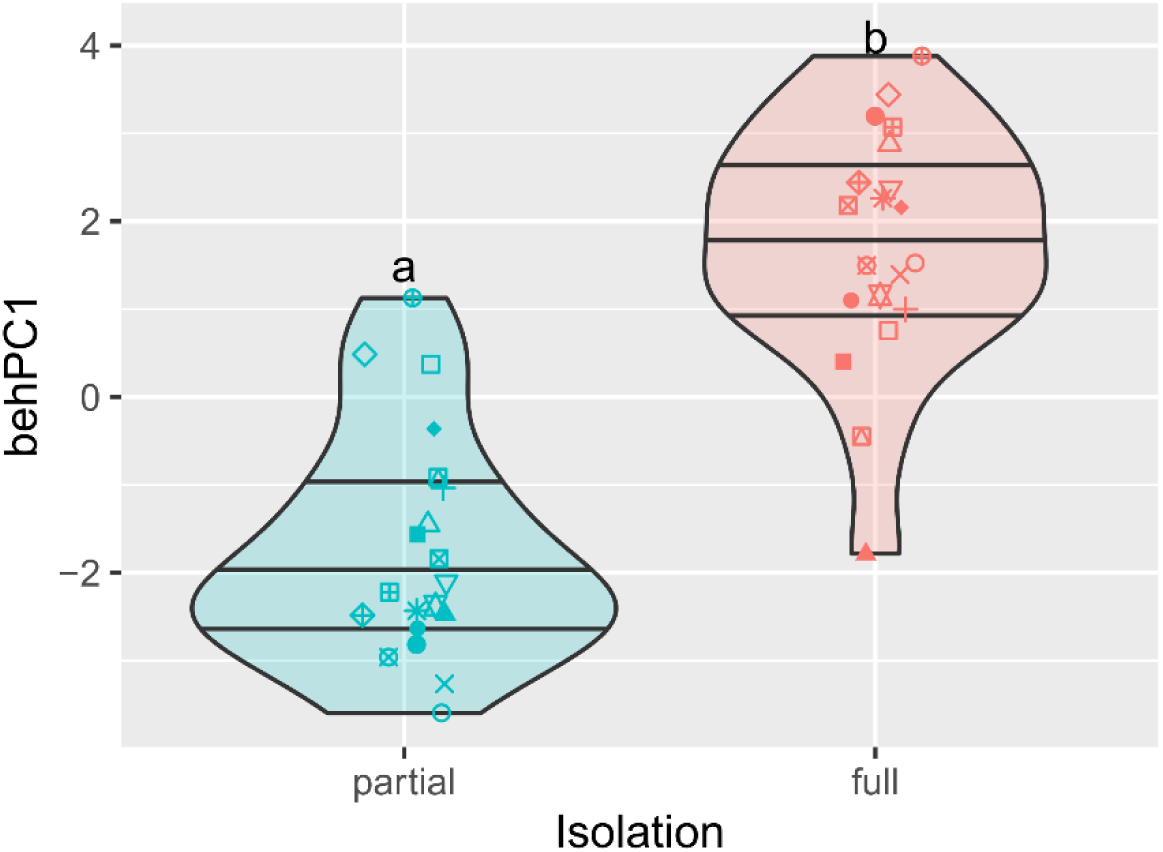
Violin plot representing the distribution of lambs’ behavioural response scores (behPC1) in partial and full isolation. Individual subject points in partial and full isolation are depicted by shapes. Horizontal lines represent first, second and third quartiles of data distribution (Anova Table and model estimates are available in supplementary Tables S2, S3, S4 respectively, model 1)

### 4.3 Thermal eye profile across partial and full isolation

The eye max temperature Coefficient of Reaction (temperatureCR) was significantly affected by the interaction between the phase of the test and lamb size (LMM main effect, Chisq = 38.6; p = <0.001; figure 2); from partial to full isolation, temperatureCR increased (i.e., became positive) in small lambs (partial *vs* full, t.ratio = 8.82, p <0.001, figure 2A), but not in large ones (partial *vs* full, t.ratio = 0.42, p = 0.68, figure 2A). In addition, the delta from the start of the phase also affected temperatureCR (LMM main effect, Chisq = 9.26; p = 0.002; figure 2B); the higher the delta, the higher the temperatureCR (slope estimate[95%CI] = 0.58 [0.21:0.96]; supplementary Tables S5-S7).

**Figure 2.**
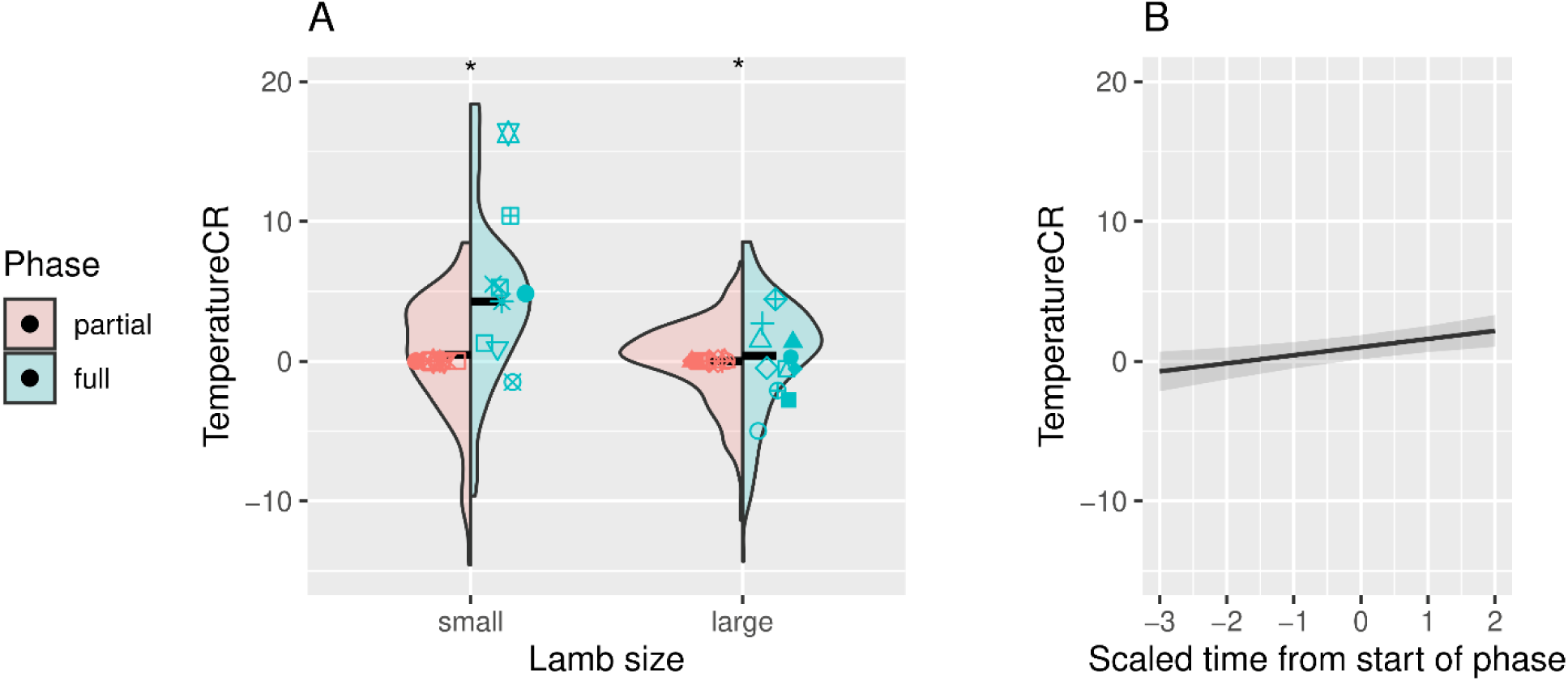
Lambs’ thermal response score (TemperatureCR) of small and large lambs in partial and full isolation (2A) and depending on the delta to the start of the phase (2B). Violin plot represents the contour of data distribution and mean and standard error of the groups are depicted with horizontal lines. Individual subject points of small and large lambs are depicted by shapes (Anova Table and model estimates are available in supplementary Tables S5 – S7).

### 4.2 Variation of high bleats acoustic structure across partial and full isolation

The Principal Component Analysis performed on the acoustic data (348 high bleats, 321 produced during the full isolation and 27 during the partial isolation) showed four components describing the spectro-temporal features of high bleats: -acPC1 to acPC4, respectively explaining 37%, 15%, 13% and 9% of the variance. According to the loadings of the acoustic features on the PCs (Table 3), higher -acPC1 scores indicated calls with a higher median, a higher Wiener entropy, a higher dominant frequency mean and sd, a higher mean of the second and third formants and a higher fundamental frequency. Therefore, as -acPC1 scores increased, calls became less tonal and were characterised by higher frequencies. Higher -acPC2 scores indicated calls with a higher second formant. Higher values of acPC3 indicated calls that were characterized by a higher value of Inter Quartile Range (higher frequency bandwidth), and finally, higher values of acPC4 indicated calls with longer duration and less variability in the fundamental frequency.

**Table 3.**
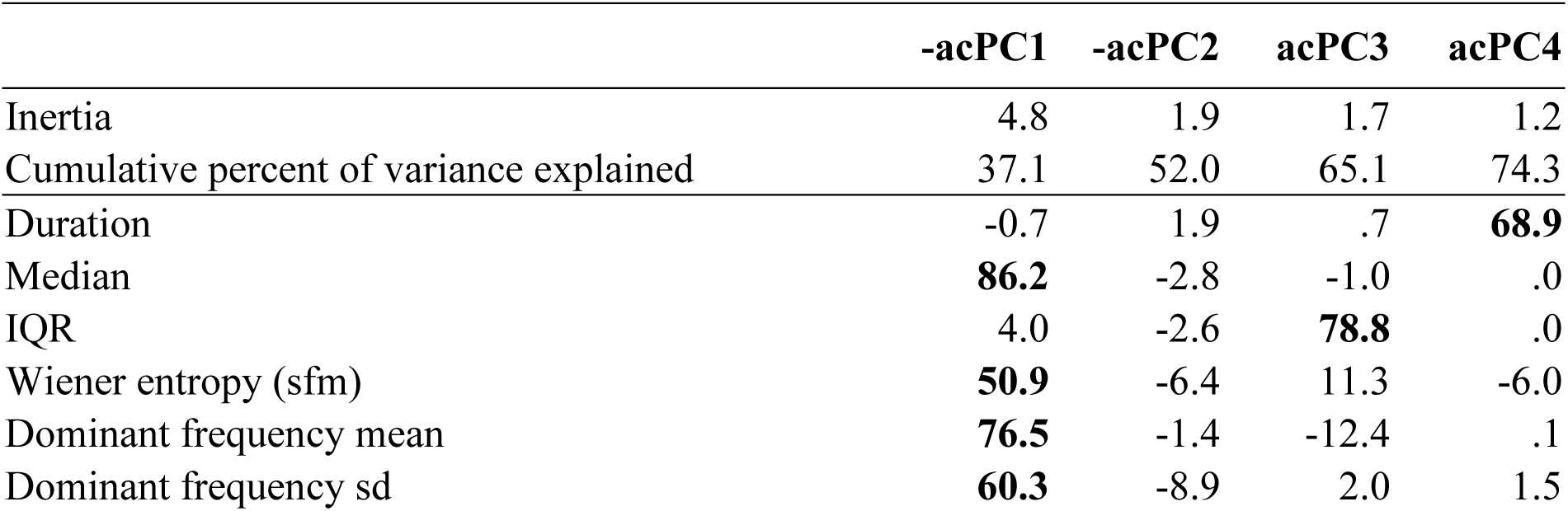
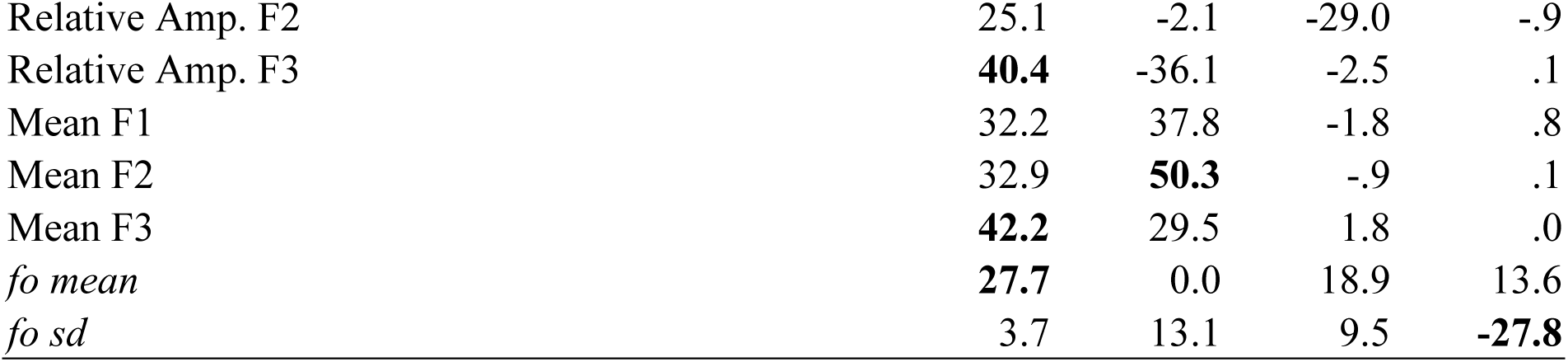
Results of the Principal Component Analysis performed on the acoustic dataset giving the features’ loadings on -acPC1,-acPC2, ac PC3 and acPC4 (relative contribution of each parameter to the PCs). The highest contribution of each acoustic feature appears in bold letters.

The LMMs showed that –acPC1 scores were significantly higher in full compared to partial isolation (LMM main effect, Chisq = 8.23; df = 1, p = 0.004, figure 3A), and were affected by the size of the lamb (LMM main effect, Chisq = 8.43; df = 1, p = 0.004, figure 3A). In addition, acPC4 scores were significantly lower in full compared to partial isolation (LMM main effect, Chisq = 11.58; df = 1, p < 0.001, figure 3C). No evidence for any effects of the phase nor the size of the lamb was found on –acPC2 (Chisq = 1.94; df = 1, p = 0.16; Chisq = 0.75; df = 1, p = 0.39, respectively) and acPC3 scores (Chisq = 2.38; df = 1, p = 0.12; Chisq = 0.45; df = 1, p = 0.50, respectively; supplementary Table S8 – S10). Hence, considering the loadings of acoustic features on the PCs (Table 3), this indicates that, compared to high bleats produced in partial isolation, bleats produced in full isolation were less tonal, composed of higher frequencies, of shorter durations and with more variability in the fundamental frequency.

**Figure 3.**
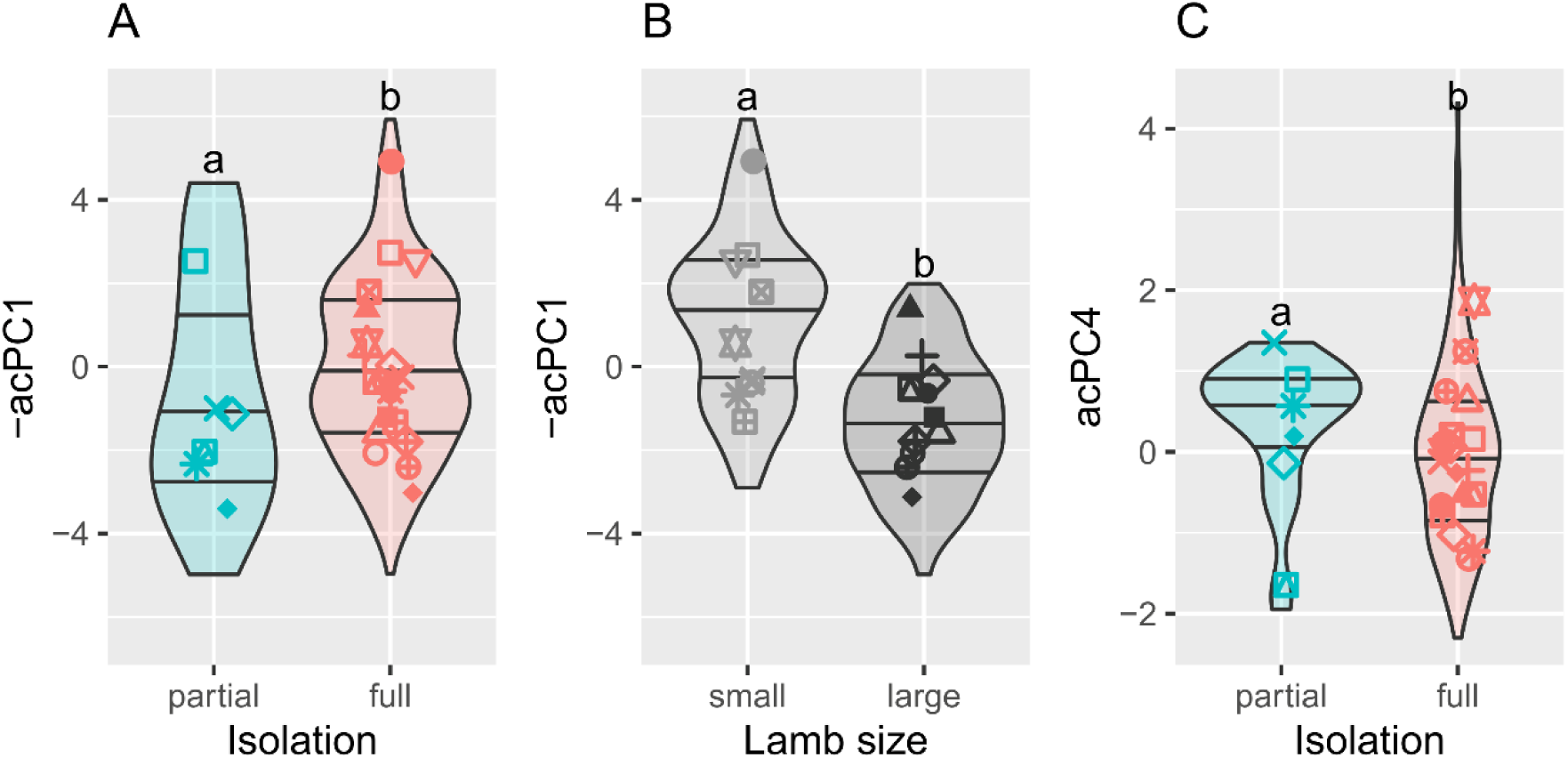
Violin plots representing the distribution of lambs’ acoustic response scores (-acPC1 and acPC4) in partial and full isolation and in small and large lambs (-acPC1). Individual subject points in partial and full isolation are depicted by shapes. Horizontal lines represent first, second and third quartiles of data distribution. (Anova Table and model estimates are available in supplementary Tables S8-S10, model 2)

### 4.4 Co-variation of acoustic, thermal and behavioural profiles

-acPC1 scores were significantly affected by behPC1: the higher the behPC1 scores, the higher – the –acPC1 scores (LMM main effect, Chisq =3.87; df = 1; p = 0.05; figure 4; Table S12). As behPC1 was found to be a descriptor of bodily activation, according to the loadings of the acoustic parameters on acoustic PCA, our results thus indicate that the higher the arousal of the lambs was during the test, the higher in frequencies and the less tonal their calls were. -acPC1 scores were significantly higher for small than for large lambs (LMM main effect, mean estimates [95% confidence interval] = 1.01[−0.21:2.23] and −1.21[−2.44:0.03], respectively Table S13). However, no evidence of any effect of the peak eye temperature was found on –acPC1 scores (LMM main effect, Chisq = 0.05 p = 0.82, Table S12). acPC4 scores were significantly affected by the interaction between behPC1 and the peak eye temperature (LMM main effect, Chisq = 7.17, p = 0.007, figure 4; Table S12). Posthoc tests investigating differences between slope trends showed that acPC4 scores significantly increased with eye peak temperature, but only for lambs having higher behPC1 scores (slope estimates [95% confidence interval] = −0.04[−0.24:0.17], 0.20[−0.08:0.49] and 0.44[0.01:0.88] respectively for low, medium and high behPC1 scores; Table S17). According to the loadings of the acoustic parameters on the acoustic PCA, this indicates that eye peak temperature during the test was positively correlated with call duration and negatively related with the variability in the fundamental frequency; however, this was true only when lambs reached a sufficiently high level of bodily activation. No effect of lamb size was found on acPC4 scores (LMM main effect, Chisq = 2.0 p = 0.16, Table S12). AcPC3 was significantly affected by the interaction between behPC1 and eye peak temperature (LMM main effect, Chisq = 4.8 p = 0.03, Table S12), however, posthoc tests did not reveal any significant slopes nor significant differences between them (Table S18). Finally, acPC3 and –acPC2 scores were significantly affected by the interaction between lamb size and eye peak temperature (LMM main effect, Chisq = 4.2 p =0.04 and Chisq = 4.0 p =0.04 respectively; Tables S12), yet, posthoc tests only revealed a slightly significant slope for –acPC2 scores, which increased with eye peak temperature in small lambs compared to large lambs (t.ratio = −2.00, p = 0.05 (slope estimates [95% confidence interval] = −0.37[−0.01:0.75], −0.09[−0.51:0.33] respectively for small and large lambs; Tables S14 and S18).

**Figure 4.**
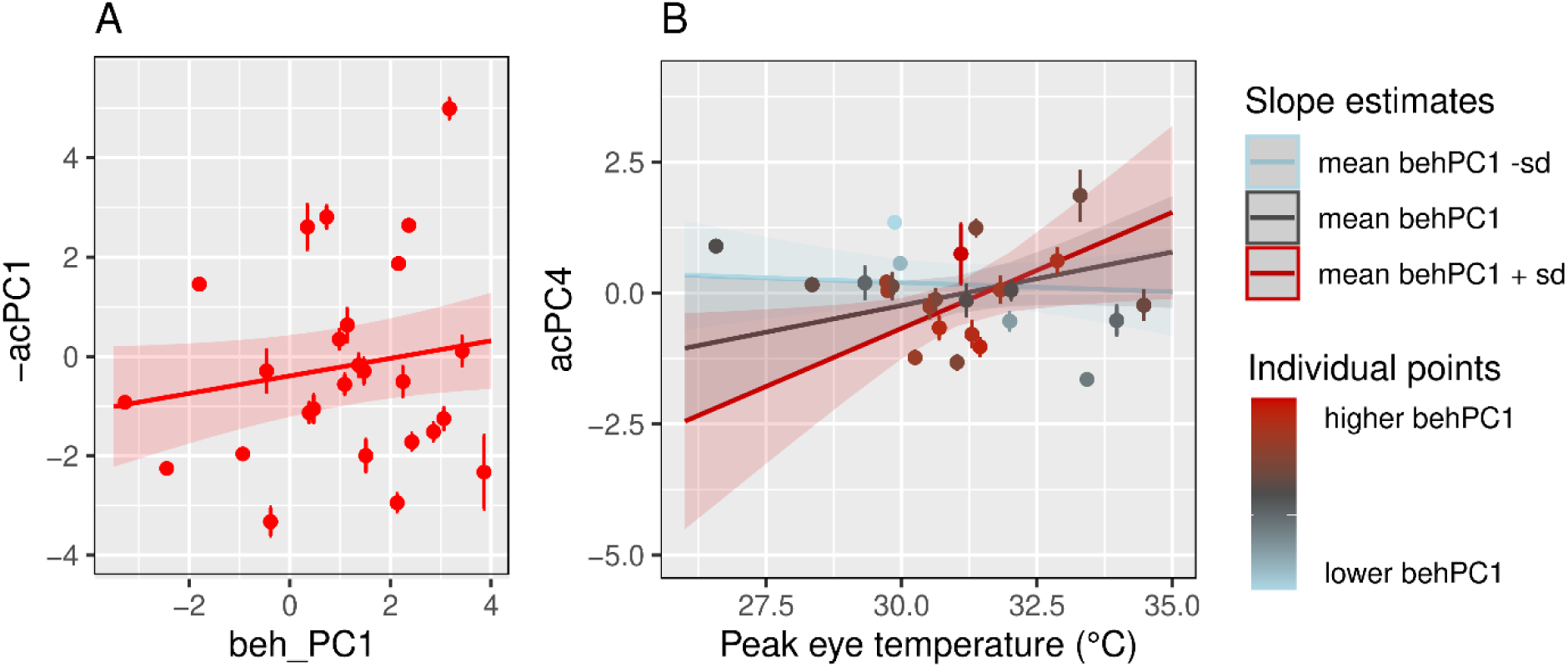
Co-variation of -acPC1 (A) and acPC4 (B) as a function of behPC1 (A) and eye peak temperature (B). Slopes depict model estimates and confidence interval. Points and segments depict individual mean and standard error. (Anova Table, model estimates and posthoc tests are available in supplementary Tables S12 – S18, model 4)

## 5. Discussion

We investigated how lambs express emotional arousal through vocalizations in response to experimentally-induced negative situations of varying intensity (i.e., partial and full isolation). Behavioural, vocal and thermal indicators differed between the two phases. When examining the relationships between bodily activation, eye peak temperature and acoustic features, we found that vocalizations’ features co-varied with bodily activation, showing higher frequencies and lower tonality when bodily activation increased. Call duration also increased, and the variability in the fundamental frequency decreased, with eye peak temperature, but only when bodily activation was sufficiently high. Below, we discuss the validity of the experimental phases, the co-variation between indicators, and their sensitivity and limits.

### 5.1 Bodily activation as a proxy for negative arousal during full isolation

During full isolation, lambs spent more time running and less time in static postures (standing and lying), performed more jumps, changes of state behaviours and bar head touches, and produced more calls than during partial isolation. These behavioural changes indicate higher arousal levels. Indeed, increased locomotor activity is commonly interpreted as a response to fear and anxiety induced by social isolation in sheep and lambs (Vandenheede et al., 1998, Engeldal et al., 2013, Ligout et al., 2011), often reflecting escape attempts (Siebert et al., 2011). Such responses are consistent with the fight-or-flight response, which is typical of highly arousing situations (Dawkins, 1998). Similar increases in locomotor activity (Briefer et al., 2015a) and escape attempts (Siebert et al., 2011) have been reported in goats exposed to high arousal conditions. However, some studies in sheep and goats have reported reduced locomotor activity during full isolation, suggesting lower arousal when communication with conspecifics is prevented (Siebert et al., 2011; Engeldal et al., 2013). These contrasting results are likely due to methodological differences, and in particular the longer duration of isolation test in those studies. Prolonged absence of contact with conspecifics may reduce reactivity, leading to behavioural attenuation or resignation.

The production of high (open-mouth) vocalisations has previously been observed in sheep exposed to stress caused by separation from offspring and social isolation (Dwyer, 2008; Papadaki et al., 2021; Hazard et al., 2022) and in lambs during social isolation and maternal separation (Ligout et al., 2011; Han et al., 2024). In our study, lambs also produced more calls per time unit (faster rate) during full isolation. This increase in vocal rate may reflect increased respiratory rate with rising emotional arousal (Briefer, 2020). Higher call rates have been observed in several species, including cattle (Thomas et al., 2001) and pigs (Marchant et al., 2001), in both negative and positive contexts (Villain et al., 2020), supporting their use as a general indicator of arousal (Briefer, 2020).

In this study, ear and tail postures could not be analysed because they were not visible for most of the time. Although some studies suggest that these postures reflect emotional states in sheep (ears: Boissy et al., 2011; Reefmann et al., 2009a; Vögeli et al., 2015; Reefmann et al., 2009a), most studies have focused on only one emotional dimension (valence or arousal), without controlling for the other. It therefore remains difficult to identify specific postures as reliable indicators of arousal, and further research is needed.

### 5.2 Increase in maximum eye temperature during full isolation

We found positive and increasing individual coefficients of reaction from partial to full isolation, indicating that maximum eye temperature rose with arousal, but only in small lambs. Several studies using infrared thermography in sheep have reported increases in eye temperature (Stubsjøen et al., 2009; Arfuso et al., 2022), or ocular region temperature (Cannas et al., 2018; Comin et al., 2024; Arfuso et al., 2022) in response to stress or fear. Here, we extracted the thermal matrices that covered the entire ocular area. The medial canthus has been identified as a reliable ocular region for monitoring stress, because it is affected by vasodilation and heat dissipation during exposure to stressors (Arfuso et al. 2022). Its temperature has also been shown to correlate positively with serum cortisol concentration and rectal temperature. At the time this study was conducted, potential lateralization effects in infrared eye temperature had not yet been investigated. Although recent evidence suggests that asymmetries may occur in calves during negative events (de Oca et al., 2024), the changes observed in our study despite not controlling for eye side suggest that the effect detected is robust.

In our study, the effect of full isolation on eye temperature depended on lamb size. Body size influences metabolic rate and thermoregulation in mammals (Kleiber, 1947). For instance, younger Standardbred trotting horses showed higher basal temperatures than adults (Negro et al., 2018). Larger lambs in our study may have had higher baseline eye temperatures, leading to a ceiling effect that limited further increases during full isolation. Ocular surface temperature also increased progressively over time, regardless of the experimental phase. This trend suggests that extracting peak eye temperatures as a proxy for arousal may be more appropriate than using average values, as averages may be influenced by temporal drift during the test.

### 5.3 Variation in vocal structure of high bleats during full isolation

Our acoustic analysis showed that lambs produced bleats with higher frequencies during full isolation, including higher formants, fundamental and dominant frequencies. Increases in vocal frequencies with rising arousal have been reported in many species (goats; Briefer et al., 2015a; piglets, Linhart et al., 2015; horses, Briefer et al., 2015b). Briefer et al. (2015a) observed increases in *fo* mean and other *fo* contour features (i.e., *fo* end) in goats experiencing high-arousal situations (i.e., feed anticipation and frustration) compared to low-arousal situations (i.e., isolation from conspecifics). In sheep, an increase in fundamental frequency has also been observed during separation from offspring (Sèbe et al., 2012). Higher *fo* may result from increased subglottal pressure caused by changes in respiration, and/or increased vocal fold tension or laryngeal muscle activity (Titze, 1994; Fant, 1960).

We also observed a shift in energy distribution towards higher frequencies with increasing arousal (median and dominance frequency), a pattern documented across species (Briefer, 2012; Briefer, 2020). In goats, such changes have been linked to reduced laryngeal retraction and/or increased pharyngeal constriction (Briefer et al., 2015a). One study in sheep reported opposite results, with reduced energy quartiles (i.e., Q25%, Q50% and Q70% under higher arousal; Sèbe et al., 2012). Engeldal et al. (2013) also found a reduction in energy distribution in vocalizations of sheep subjected to partial compared to full isolation (however, differences in experimental protocols, particularly longer isolation duration, may explain these discrepancies, as prolonged isolation may reduce arousal over time). Finally, higher formant frequencies have been predicted as a result of increased pharyngeal constriction and tension of the vocal tract walls as well as reduced salivation in humans (Scherer, 1986), which could explain our findings in sheep, considering similarities in vocal production in mammals (mean F3).

We also found that lamb calls became less tonal (i.e. more noisy or chaotic, as shown by an increased Wiener entropy (sfm)), as arousal rose. This finding aligns with Siebert et al. (2011), who reported more tonal calls in goat calls during 30min complete isolation, corresponding to reduced arousal probably from prolonged absence of sensory contact and social motivation. In pigs, chaos increased during anticipation regardless of valence (Villain et al., 2020), although how tonality/chaos varies with arousal may depend on the type of call (Linhart et al., 2015). An increase in chaos with arousal has been reported across species, as shown also in dog puppies (Massenet et al., 2025), marmots (Blumstein, 2025), and bonobos (Pisanski et al., 2022).

The acoustic PCA also suggested that call duration decreased from partial to full isolation. Across species, call duration is often linked to emotional valence rather than arousal (Briefer, 2020), with longer calls typically produced in negative contexts (Sèbe et al., 2007; Briefer, 2020; Villain et al., 2020; Lefèvre et al., 2025). However, duration may also increase or decrease with arousal depending on the call type. Alarm calls, for example, may become shorter at higher arousal because they must be produced rapidly (Briefer, 2012; Zimmermann et al. 2013). In our study, higher arousal during full isolation may have favoured shorter calls that could be repeated quickly, as indicated by the increased call rate. This suggests a trade-off between call duration and production rate, where shorter calls allow faster signalling under highly arousing conditions.

### 5.4 Relationship between vocal, behavioural and physiological indicators

Our combined analysis showed that acoustic features were primarily modulated by bodily activation and, to a lesser extent, by the interaction with body size and physiological arousal. Higher bodily activation was associated with increased frequencies and reduced tonality, indicating more chaotic vocal structure. Peak eye temperature alone was not directly related with these acoustic features, but it interacted with bodily activation to influence call duration. Call duration increased with eye temperature only in highly activated lambs. This suggests that physiological arousal may affect vocal output primarily when behavioural arousal is already elevated. However, the direction of change in call duration differed between analyses focusing solely on vocal data and those combining multiple indicators. This discrepancy may be due to the small number of lambs vocalising during partial isolation, which may have influenced the results. We also observed a positive relationship between the acoustic score reflecting mean of the second formant and eye peak temperature, depending on lamb size. This trend was present in small lambs but not in larger ones, supporting the hypothesis of a ceiling effect in larger individuals. These findings highlight the importance of considering biometric factors when validating thermographic indicators. Overall, these findings highlight that vocal expression of emotional arousal in lambs reflects a complex interplay between behavioural and physiological indicators, with potential modulation by body size. Vocalizations therefore appear to provide a robust and accessible proxy for emotional arousal in lambs exposed to negative situations.

## 6. Conclusion

This study revealed promising non-invasive vocal indicators of negative arousal in lambs. We found that vocalizations were reliable indicators of negative arousal levels in lambs and were correlated with bodily activation, with higher frequencies and decreased tonality (more chaos) observed as arousal increased. Co-variations with physiological measures, however, depended on both behavioural activation and the animal’s biometric characteristics. Our study provides a methodology to study the interplay between behavioural, physiological and vocal responses to emotional arousal. These findings highlight the need for a multi-disciplinary approach to accurately assess emotional arousal in animals.

## 7. Acknowledgements

We are grateful to Selma Bergenholz from Højbogaard farm for her willingness to participate in this project, but also for sharing her experience and knowledge.

## Bibliography

Arfuso, F., Acri, G., Piccione, G., Sansotta, C., Fazio, F., Giudice, E., & Giannetto, C. (2022). Eye surface infrared thermography usefulness as a noninvasive method of measuring stress response in sheep during shearing: Correlations with serum cortisol and rectal temperature values. Physiology and Behavior, 250. 10.1016/j.physbeh.2022.113781

Baciadonna, L., Nawroth, C., & McElligott, A. G. (2016). Judgement bias in goats (Capra hircus): Investigating the effects of human grooming. PeerJ, 2016(10). 10.7717/peerj.2485

Bates, D., Mächler, M., Bolker, B. M., & Walker, S. C. (2015). Fitting linear mixed-effects models using lme4. Journal of Statistical Software, 67(1). 10.18637/jss.v067.i01

Blumstein, D. T. (2025). Nonlinear phenomena in marmot alarm calls: A mechanism encoding fear? In Philosophical Transactions of the Royal Society B: Biological Sciences (Vol. 380, Issue 1923). Royal Society Publishing. 10.1098/rstb.2024.0008

Boissy, A., Aubert, A., Désiré, L., Greiveldinger, L., Delval, E., & Veissier, I. (2011). Cognitive sciences to relate ear postures to emotions in sheep. Animal Welfare, 20(1), 47–56. 10.1017/s0962728600002426

Bradley, M. M., Codispoti, M., Cuthbert, B. N., & Lang, P. J. (2001). Emotion and Motivation I: Defensive and Appetitive Reactions in Picture Processing. Emotion, 1(3), 276–298. 10.1037/1528-3542.1.3.276

Briefer, E. F. (2012). Vocal expression of emotions in mammals: Mechanisms of production and evidence. In Journal of Zoology (Vol. 288, Issue 1, pp. 1–20). 10.1111/j.1469-7998.2012.00920.x

Briefer, E. F. (2020). Coding for ‘Dynamic’ Information: Vocal Expression of Emotional Arousal and Valence in Non-human Animals (pp. 137–162). 10.1007/978-3-030-39200-0_6

Briefer, E. F., Maigrot, A. L., Mandel, R., Freymond, S. B., Bachmann, I., & Hillmann, E. (2015b). Segregation of information about emotional arousal and valence in horse whinnies. Scientific Reports, 4. 10.1038/srep09989

Briefer, E. F., Tettamanti, F., & McElligott, A. G. (2015a). Emotions in goats: Mapping physiological, behavioural and vocal profiles. Animal Behaviour, 99, 131–143. 10.1016/j.anbehav.2014.11.002

Cannas, S., Palestrini, C., Canali, E., Cozzi, B., Ferri, N., Heinzl, E., Minero, M., Chincarini, M., Vignola, G., & Dalla Costa, E. (2018). Thermography as a non-invasive measure of stress and fear of humans in sheep. Animals, 8(9). 10.3390/ani8090146

Cockram, M. S., Ranson, M., Imlah, P., Goddard, P. J., Burrells, C., & Harkiss, G. D. (1994). The behavioural, endocrine and immune responses of sheep to isolation. Animal Science, 58(3), 389–399.

Comin, M., Atallah, E., Chincarini, M., Mazzola, S. M., Canali, E., Minero, M., Cozzi, B., Rossi, E., Vignola, G., & Dalla Costa, E. (2024). Events with Different Emotional Valence Affect the Eye’s Lacrimal Caruncle Temperature Changes in Sheep. Animals, 14(1). 10.3390/ani14010050

Davies, A. C., Radford, A. N., & Nicol, C. J. (2014). Behavioural and physiological expression of arousal during decision-making in laying hens. Physiology and Behavior, 123, 93–99. 10.1016/j.physbeh.2013.10.008

Dawkins, M. S. (1998). Evolution and animal welfare. Quarterly Review of Biology, 73(3), 305–328. 10.1086/420307

de Oca, M. A. R. M., Mendl, M., Whay, H. R., Held, S. D., Lambton, S. L., & Telkänranta, H. (2024). An exploration of surface temperature asymmetries as potential markers of affective states in calves experiencing or observing disbudding. Animal Welfare, 33, e45. 10.1017/awf.2024.47

de Oliveira, D., & Keeling, L. J. (2018). Routine activities and emotion in the life of dairy cows: Integrating body language into an affective state framework. PLoS ONE, 13(5). 10.1371/journal.pone.0195674

Dray, S., & Dufour, A.-B. (2007). Journal of Statistical Software The ade4 Package: Implementing the Duality Diagram for Ecologists. http://www.jstatsoft.org/

Dwyer, C. M. (2008). The welfare of sheep (Springer, Ed.). Springer Dordrecht. 10.1007/978-1-4020-8553-6

Engeldal, S. E. C., Handiwirawan, E., & Noor, R. R. (2013). Effect of different levels of social isolation on the acoustical characteristics of sheep vocalization. Jurnal Ilmu Ternak Dan Veteriner, 18(3), 208–219.

Ermatinger, F. A., Brügger, R. K., & Burkart, J. M. (2019). The use of infrared thermography to investigate emotions in common marmosets. Physiology and Behavior, 211. 10.1016/j.physbeh.2019.112672

Fant, G. (1960). FANT, Gunnar. Acoustic theory of speech production (Mouton, Ed.).

Fox, J., & Weisberg, S. (2018). An R companion to applied regression (Sage publications, Ed.).

Friard, O., & Gamba, M. (2016). BORIS: a free, versatile open-source event-logging software for video/audio coding and live observations. Methods in Ecology and Evolution, 7(11), 1325–1330. 10.1111/2041-210X.12584

Han, C., Li, M., Li, F., Wang, Z., Hu, X., Yang, Y., Wang, H., & Lv, S. (2024). Temporary sensory separation of lamb groups from ewes affects behaviors and serum levels of stress-related indicators of small-tailed Han lambs. Physiology & Behavior, 277, 114504. 10.1016/j.physbeh.2024.114504

Ingram, J. R., Cook, C. J., & Harris, P. J. (2002). The Effect of Transport on Core and Peripheral Body Temperatures and Heart Rate of Sheep. Animal Welfare, 11(1), 103–112.

Kleiber, M. (1947). Body size and metabolic rate. Physiological Reviews, 27(4), 511–541.

Lefèvre, R. A., Sypherd, C. C. R., & Briefer, É. F. (2025). Machine learning algorithms can predict emotional valence across ungulate vocalizations. IScience, 28(2). 10.1016/j.isci.2025.111834

Lenth, R. V. (2016). Least-squares means: The R package lsmeans. Journal of Statistical Software, 69. 10.18637/jss.v069.i01

Ligges, U., Krey, S., Mersmann, O., Schnackenberg, S., Guénard, G., Ellis, D. P. W., & Underbit Technologies. (2023). tuneR: Analysis of Music and Speech.

Ligout, S., Foulquié, D., Sèbe, F., Bouix, J., & Boissy, A. (2011). Assessment of sociability in farm animals: The use of arena test in lambs. Applied Animal Behaviour Science, 135(1–2), 57–62. 10.1016/j.applanim.2011.09.004

Linhart, P., Ratcliffe, V. F., Reby, D., & Špinka, M. (2015). Expression of emotional arousal in two different piglet call types. PLoS ONE, 10(8). 10.1371/journal.pone.0135414

Lowe, T. E., Cook, C. J., Ingram, J. R., & Harris, P. J. (2005). Changes in ear-pinna temperature as a useful measure of stress in sheep (Ovis aries). Animal Welfare, 14(1), 35–42.

Marcet Rius, M., Pageat, P., Bienboire-Frosini, C., Teruel, E., Monneret, P., Leclercq, J., Lafont-Lecuelle, C., & Cozzi, A. (2018). Tail and ear movements as possible indicators of emotions in pigs. Applied Animal Behaviour Science, 205, 14–18. 10.1016/j.applanim.2018.05.012

Marchant, J. N., Whittaker, X., & Broom, D. M. (2001). Vocalisations of the adult female domestic pig during a standard human approach test and their relationships with behavioural and heart rate measures. Applied Animal Behaviour Science, 72(1), 23–29.

Mason, M. (2023). Goat Perception of Human Cues & Reliability of Thermal Imaging in Welfare Research.

Massenet, M., Mathevon, N., Anikin, A., Briefer, E. F., Fitch, W. T., & Reby, D. (2025). Nonlinear phenomena in vertebrate vocalizations: Mechanisms and communicative functions. In Philosophical Transactions of the Royal Society B: Biological Sciences (Vol. 380, Issue 1923). Royal Society Publishing. 10.1098/rstb.2024.0002

Mendl, M., Burman, O. H. P., & Paul, E. S. (2010). An integrative and functional framework for the study of animal emotion and mood. Proceedings of the Royal Society B: Biological Sciences, 277(1696), 2895–2904. 10.1098/rspb.2010.0303

Mendl, M., & Paul, E. S. (2020). Animal affect and decision-making. In Neuroscience and Biobehavioral Reviews (Vol. 112, pp. 144–163). Elsevier Ltd. 10.1016/j.neubiorev.2020.01.025

Negro, S., Bartolomé, E., Molina, A., Solé, M., Gómez, M. D., & Valera, M. (2018). Stress level effects on sport performance during trotting races in Spanish Trotter Horses. Research in Veterinary Science, 118, 86–90. 10.1016/j.rvsc.2018.01.017

Papadaki, K., Laliotis, G. P., & Bizelis, I. (2021). Acoustic variables of high-pitched vocalizations in dairy sheep breeds. Applied Animal Behaviour Science, 241. 10.1016/j.applanim.2021.105398

Pisanski, K., Bryant, G. A., Cornec, C., Anikin, A., & Reby, D. (2022). Form follows function in human nonverbal vocalisations. In Ethology Ecology and Evolution (Vol. 34, Issue 3, pp. 303–321). Taylor and Francis Ltd. 10.1080/03949370.2022.2026482

Proctor, H. S., & Carder, G. (2015). Nasal temperatures in dairy cows are influenced by positive emotional state. Physiology and Behavior, 138, 340–344. 10.1016/j.physbeh.2014.11.011

Reefmann, N., Bütikofer Kaszàs, F., Wechsler, B., & Gygax, L. (2009a). Ear and tail postures as indicators of emotional valence in sheep. Applied Animal Behaviour Science, 118(3–4), 199–207. 10.1016/j.applanim.2009.02.013

Reefmann, N., Wechsler, B., & Gygax, L. (2009b). Behavioural and physiological assessment of positive and negative emotion in sheep. Animal Behaviour, 78(3), 651–659. 10.1016/j.anbehav.2009.06.015

Rigterink, A., Moore, G. E., & Ogata, N. (2018). Pilot study evaluating surface temperature in dogs with or without fear-based aggression. Journal of Veterinary Behavior, 28, 11–16. 10.1016/j.jveb.2018.07.009

Russell, J. A. (1980). A circumplex model of affect. Journal of Personality and Social Psychology, 39(6), 1161.

Scherer, K. R. (1986). Vocal affect expression: a review and a model for future research. Psychological Bulletin, 99(2), 143–165.

Sèbe, F., Nowak, R., Poindron, P., & Aubin, T. (2007). Establishment of vocal communication and discrimination between ewes and their lamb in the first two days after parturition. Developmental Psychobiology, 49(4), 375–386. 10.1002/dev.20218

Sèbe, F., Poindron, P., Andanson, S., Chandeze, H., Delval, E., Despres, G., Toporenki, G., Bickell, S., & Boissy, A. (2012). Bio-acoustic analyses to assess emotion in animals: acoustic patters are linked to behavioural, cardiac and hormonal responses of ewes to the separation from their lambs. In Biacoustics (Ed.), Proceedings of the International Bioacoustics Council meeting (p. 54). Biacoustics.

Siebert, K., Langbein, J., Schön, P. C., Tuchscherer, A., & Puppe, B. (2011). Degree of social isolation affects behavioural and vocal response patterns in dwarf goats (Capra hircus). Applied Animal Behaviour Science, 131(1–2), 53–62. 10.1016/j.applanim.2011.01.003

Stubsjøen, S. M., Flø, A. S., Moe, R. O., Janczak, A. M., Skjerve, E., Valle, P. S., & Zanella, A. J. (2009). Exploring non-invasive methods to assess pain in sheep. Physiology and Behavior, 98(5), 640–648. 10.1016/j.physbeh.2009.09.019

Sueur, J., Aubin, T., & Simonis, C. (2008). Seewave, a free modular tool for sound analysis and synthesis. Bioacoustics, 18(2), 213–226. http://cran.r-project.org/src/contrib/Descriptions/

Taylor, A. M., & Reby, D. (2010). The contribution of source-filter theory to mammal vocal communication research. Journal of Zoology, 280(3), 221–236. 10.1111/j.1469-7998.2009.00661.x

Thomas, T. J., Weary, D. M., & Appleby, M. C. (2001). Newborn and 5-week-old calves vocalize in response to milk deprivation. Applied Animal Behaviour Science, 74(3), 165–173.

Titze, I. R. (1994). Principles of voice production.

Travain, T., & Valsecchi, P. (2021). Infrared thermography in the study of animals’ emotional responses: A critical review. In Animals (Vol. 11, Issue 9). MDPI AG. 10.3390/ani11092510

Vandenheede, M., Bouissou, M. F., & Picard, M. (1998). Interpretation of behavioural reactions of sheep towards fear-eliciting situations. In Applied Animal Behaviour Science (Vol. 58).

Vianna, D. M. L., & Carrive, P. (2005). Changes in cutaneous and body temperature during and after conditioned fear to context in the rat. European Journal of Neuroscience, 21(9), 2505–2512. 10.1111/j.1460-9568.2005.04073.x

Villain, A., & Renaud-Goud, P. (2023). SoundChunk R Package and example data.

Villain, A. S., & Briefer, E. F. (n.d.). Vocal signals as indicators of emotions. In Assessment of Animal Welfare - a guide to the valid use of indicators of affective states. John Wiley & Sons Ltd.

Villain, A. S., Hazard, A., Danglot, M., Guérin, C., Boissy, A., & Tallet, C. (2020). Piglets vocally express the anticipation of pseudo-social contexts in their grunts. Scientific Reports, 10(1). 10.1038/s41598-020-75378-x

Vögeli, S., Wolf, M., Wechsler, B., & Gygax, L. (2015). Housing conditions influence cortical and behavioural reactions of sheep in response to videos showing social interactions of different valence. Behavioural Brain Research, 284, 69–76. 10.1016/j.bbr.2015.02.007

Webster, M. M., & Rutz, C. (2020). How STRANGE are your study animals? Nature, 582(7812), 337–340.

Zimmermann, E., Leliveld, L., & Schehka, S. (2013). Towards the evolutionary roots of affective prosody in human acoustic communication: a com-parative approach to mammalian voices. In Oxford University Press (Ed.), Altenmüller E, Schmidt S, Zimmermann E (eds) Evolution of emotional communication: from sounds in nonhuman mammals to speech and music in man (pp. 116–132). Oxford University Press.

