## Supplementary materials for "Acoustic markers of negative arousal in lambs: evidence from behavioural and eye thermal profiles"

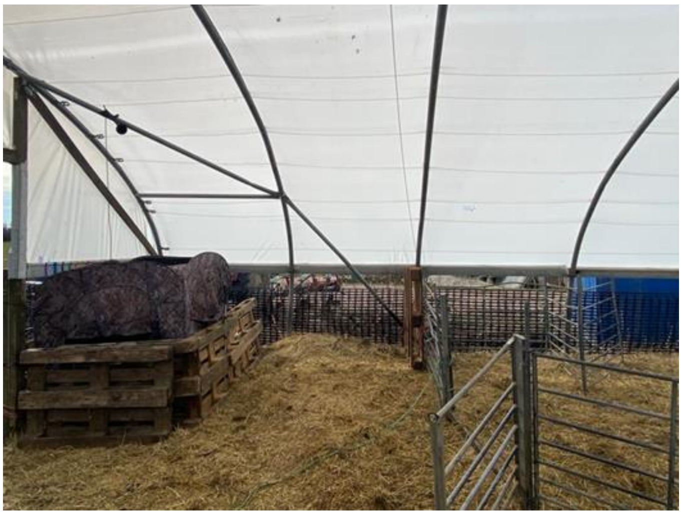

*Figure S1. Experimenter hide (top left) and the isolation pen (top right)*

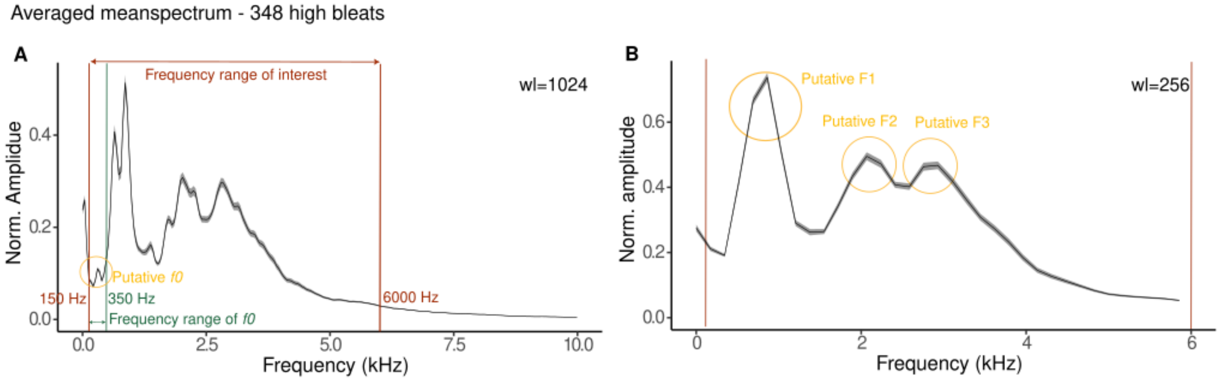


*Figure S2: Spectral representation of High Bleats, prior to acoustic analysis. Averaged meanspectrun (solid line) and standard error (shade), over the total amount of vocalisations (recorded using frequency sampling 44.1 kHz and 16 Bit). A: using a window length of 1024 (used of later analyses) allow to visualise the frequency range of interest (150-6000 Hz) used for bandpass filtering and frequency range of the fundamental frequency (f0) (150-350 Hz). B: using a window length of 256 to identify the number and position of putative formant that should be included in the analysis.*

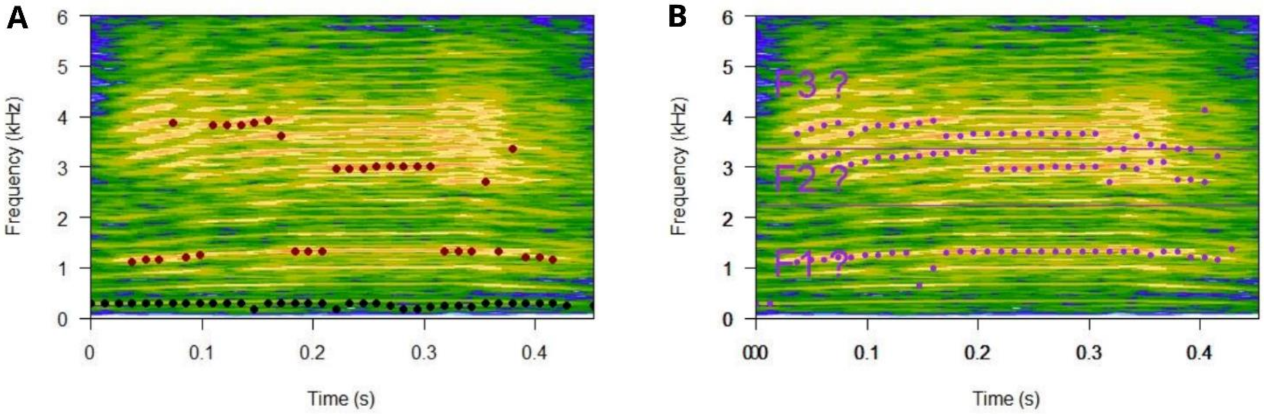


*Figure S3A. Spectrogram of a high bleat (computed with FFT wl = 2048 and ov = 90) with illustration of detection of dominant frequencies (red dots) and fundamental frequency (fo;* black dots), both *computed over the duration of the call in each sliding time window of the FFT computation (wl =1024 and ov = 50%, using either the frequency range of interest (150-6000Hz) or the frequency range expected for fo (150-350Hz) respectively for the dominant frequencies and the fundamental frequencies. Figure S3B. Spectrogram of a high bleat (computed with FFT wl = 2048 and ov = 90) with illustration of the three main putative formant frequency bands (F1?, F2? and F3?) between 350 and 6000 kHz, within which the dominant frequency of each formant was computed (purple dots for F1, F2, and F3 respectively).*

*Table S1. Acoustic features extracted from the high bleats and used for acoustic analysis*

| Acoustic feature | Unit | Description |
|---|---|---|
| Duration | s | Total duration of the vocalisation |
| Median | Hz | Frequency below which 50 % of the energy is contained (also termed 'Q50%' or 'spectral centre of gravity') |
| Inter Quartile Range (IQR) | Hz | Frequency range between the third and the first quartile (indicates the bandwidth) |
| Wiener entropy (sfm) | - | Ratio of the geometric mean to the arithmetic mean of the spectrum (0: pure tone; 1: random noise) |
| Dominant frequency mean | Hz | Mean of the dominant frequency, computed per FFT window over the duration of the vocalization |
| Dominant frequency sd | Hz | Standard deviation of the dominant frequency, computed per FFT window over the duration of the vocalization |
| Relative Amp. F2* | dB | Relative amplitude of the second formant |
| Relative Amp. F3* | dB | Relative amplitude of the third formant |
| Mean F1 | Hz | Mean frequency value of the first formant |
| Mean F2 | Hz | Mean frequency value of the second formant |
| Mean F3 | Hz | Mean frequency value of the third formant |
| $f_o$ mean | Hz | Mean of the fundamental frequency of the vocalization over the duration of the vocalisation, computed per FFT window over the duration of the vocalization |
| $f_o$ sd | Hz | Standard deviation of the fundamental frequency of the vocalization, computed per FFT window over the duration of the vocalization |

** relative amplitude= normalized amplitude (the amplitude of the bleats has been reduced or scaled so that they fall between 0 and 1; for each sound, the loudest amplitude has been set to 1 and the other sounds have been normalised between 0 and 1)*

### Statistical tables

### Tables for model 1.

*Table S2. Linear Mixed-Effects Model results (referred to behaviour PC1 and PC2)*

| Response variable | Fixed effect | Chisq | Df | Pr(>Chisq) |
|---|---|---|---|---|
| BehPC1 | Phase of the test | 96.32 | 1 | < 0.0001*** |

| | | | | |
|---|---|---|---|---|
| | Lamb size | 0.13 | 1 | 0.71 |
| | Phase of the test*lamb size | 3.41 | 1 | 0.06 |
| BehPC2 | Phase of the test | 2.34 | 1 | 0.13 |
| | Lamb size | 2.92 | 1 | 0.09 |
| | Phase of the test*lamb size | 0.13 | 1 | 0.72 |

*Significance codes: * = 0.05; ** = 0.01; *** = 0.001*

*Table S3. Model estimates (referred to behaviour PC1)*

| Phase of the test | emmean | SE | df | lower.CL | upper.CL |
|---|---|---|---|---|---|
| Full isolation | 1.75 | 0.298 | 33 | 1.14 | 2.36 |
| Partial isolation | -1.77 | 0.298 | 33 | -2.37 | -1.16 |

*Table S4. Post hoc test (referred to PC1)*

| Contrast | estimate | SE | df | t.ratio | p.value |
|---|---|---|---|---|---|
| Full isolation – partial isolation | 3.52 | 0.35 | 18 | 9.95 | <0.0001 |

### Tables for model 2.

*Table S5. Linear Mixed-Effects Model results results (referred to the Coefficient of Reaction)*

| Fixed effect | Chisq | Df | Pr(>Chisq) |
|---|---|---|---|
| Phase of the test | 41.3 | 1 | < 0.0001*** |
| Lamb size | 2.49 | 1 | 0.11 |
| Delta from phase start | 9.26 | 1 | 0.002** |
| Phase of the test*lamb size | 38.6 | | <0.0001*** |
| Phase of the test * delta from phase start | 0.18 | 1 | 0.67 |

*Significance codes: * = 0.05; ** = 0.01; *** = 0.001*

*Table S6. Model estimates (referred to the Coefficient of Reaction)*

| Phase of the test | Lamb size | emmean | SE | df | lower.CL | upper.CL |
|---|---|---|---|---|---|---|
| Full isolation | Large | 0.28 | 0.65 | 25.5 | -1.05 | 1.61 |
| Partial isolation | Large | 0.07 | 0.63 | 21.5 | -1.23 | 1.37 |
| Full isolation | Small | 4.21 | 0.73 | 25.8 | 2.72 | 5.70 |

| | | | | | | |
|---|---|---|---|---|---|---|
| Partial isolation | Small | -0.54 | 0.70 | 21.8 | -1.99 | 0.92 |

*Table S7. Post hoc test (referred to the Coefficient of Reaction)*

| Contrast | estimate | SE | df | t.ratio | p.value |
|---|---|---|---|---|---|
| Full large – partial large | 0.21 | 0.51 | 720 | 0.42 | 0.68 |
| Full small – partial small | 4.75 | 0.54 | 730 | 8.82 | <0.0001 |

### Tables for model 3 (all the lambs n = 20).

*Table S8. Linear Mixed-Effects Model results (referred to acoustic PC1, PC2, PC3 and PC4)*

| Response variable | Fixed effect | Chisq | Df | Pr(>Chisq) |
|---|---|---|---|---|
| AcPC1 | Phase of the test | 8.23 | 1 | 0.004** |
| | Lamb size | 8.43 | 1 | 0.003** |
| | Phase of the test*lamb size | 0.63 | 1 | 0.43 |
| AcPC2 | Phase of the test | 1.94 | 1 | 0.16 |
| | Lamb size | 0.75 | 1 | 0.39 |
| | Phase of the test*lamb size | 0.74 | 1 | 0.39 |
| AcPC3 | Phase of the test | 2.38 | 1 | 0.12 |
| | Lamb size | 0.45 | 1 | 0.50 |
| | Phase of the test*lamb size | 2.68 | 1 | 0.10 |
| AcPC4 | Phase of the test | 11.6 | 1 | < 0.0001*** |
| | Lamb size | 0.85 | 1 | 0.36 |
| | Phase of the test*lamb size | 1.56 | 1 | 0.21 |

*Significance codes: * = 0.05; ** = 0.01; *** = 0.001*

*Table S9. Models estimates (referred to acoustic PC1 and PC4)*

| Response variable | Fixed effect | | emmean | SE | df | lower.CL | upper.CL |
|---|---|---|---|---|---|---|---|
| AcPC1 | Phase of the test | Full isolation | 0.03 | 0.37 | 18.0 | -0.76 | 0.82 |
| | | Partial isolation | -0.54 | 0.42 | 28.7 | -1.40 | 0.32 |
| | Lamb size | Large | -1.42 | 0.52 | 20.3 | -2.49 | -0.34 |

| | | | | | | | |
|---|---|---|---|---|---|---|---|
| | | Small | 0.90 | 0.57 | 20.1 | -0.29 | 2.09 |
| AcPC4 | Phase of the test | Full isolation | -0.07 | 0.18 | 17.9 | -0.44 | 0.31 |
| | | Partial isolation | -0.61 | 0.25 | 68.6 | 0.10 | 1.11 |

*Table S10. Models estimates (referred to acoustic PC1 and PC4)*

| Response variable | Contrast | estimate | SE | df | t.ratio | p.value |
|---|---|---|---|---|---|---|
| AcPC1 | Full isolation – partial isolation | 0.57 | 0.20 | 328 | 2.78 | 0.01 |
| | Large - small | -2.32 | 0.77 | 20.1 | -3.01 | 0.01 |
| AcPC4 | Full large – partial large | -0.67 | 0.19 | 333 | -3.50 | 0.001 |

**Table for model 3 (subset – only 6 lambs that produced calls in both phases).**

*Table S11. Linear Mixed-Effects Model results (referred to acoustic PC1, PC2, PC3 and PC4)*

| Response variable | Fixed effect | Chisq | Df | Pr(>Chisq) |
|---|---|---|---|---|
| AcPC1 | Phase of the test | 5.95 | 1 | 0.015* |
| | Lamb size | 1.73 | 1 | 0.19 |
| | Phase of the test*lamb size | 0.48 | 1 | 0.49 |
| AcPC2 | Phase of the test | 1.06 | 1 | 0.30 |
| | Lamb size | 0.09 | 1 | 0.76 |
| | Phase of the test*lamb size | 0.58 | 1 | 0.44 |
| AcPC3 | Phase of the test | 2.13 | 1 | 0.14 |
| | Lamb size | 0.00 | 1 | 1.00 |
| | Phase of the test*lamb size | 3.06 | 1 | 0.08 |
| AcPC4 | Phase of the test | 16.4 | 1 | < 0.0001*** |
| | Lamb size | 0.48 | 1 | 0.49 |
| | Phase of the test*lamb size | 2.06 | 1 | 0.15 |

**Tables for model 4**

*Table S12. Linear Mixed-Effects Model results (referred to acoustic PC1, PC2, PC3 and PC4)*

| Response variable | Fixed effect | Chisq | Df | Pr(>Chisq) |
|---|---|---|---|---|
| AcPC1 | Behavioral score | 3.87 | 1 | 0.05* |

| | | | | |
|---|---|---|---|---|
| | Eye peak temperature | 0.05 | | 0.82 |
| | Lamb size | 6.45 | 1 | 0.01* |
| | Behavioral score*eye peak temperature | 0.13 | 1 | 0.72 |
| | Eye peak temperature*lamb size | 0.62 | 1 | 0.43 |
| AcPC2 | Behavioral score | 2.30 | 1 | 0.13 |
| | Eye peak temperature | 0.53 | | 0.47 |
| | Lamb size | 1.23 | 1 | 0.27 |
| | Behavioral score*eye peak temperature | 0.60 | 1 | 0.44 |
| | Eye peak temperature*lamb size | 4.24 | 1 | 0.04* |
| AcPC3 | Behavioral score | 2.83 | 1 | 0.09 |
| | Eye peak temperature | 4.95 | | 0.03* |
| | Lamb size | 0.00 | 1 | 0.99 |
| | Behavioral score*eye peak temperature | 4.81 | 1 | 0.03* |
| | Eye peak temperature*lamb size | 4.02 | 1 | 0.04* |
| AcPC4 | Behavioral score | 7.96 | 1 | 0.005** |
| | Eye peak temperature | 0.20 | | 0.65 |
| | Lamb size | 1.98 | 1 | 0.16 |
| | Behavioral score*eye peak temperature | 7.18 | 1 | 0.007** |
| | Eye peak temperature*lamb size | 0.05 | 1 | 0.83 |

*Significance codes: * = 0.05; ** = 0.01; *** = 0.001*

*Table S13. Model estimates (referred to acoustic PC1)*

| **Response variable** | **Lamb size** | **emmean** | **SE** | **df** | **lower.CL** | **upper.CL** |
|---|---|---|---|---|---|---|
| AcPC1 | Large | -1.21 | 0.59 | 19.8 | -2.44 | 0.03 |
| | Small | 1.01 | 0.58 | 16.5 | -0.21 | 2.23 |

*Table S14. Model estimates (referred to acoustic PC2)*

| Lamb size | Eye peak temperature | SE | df | lower.CL | upper.CL |
|---|---|---|---|---|---|
| Large | -0.09 | 0.20 | 19.2 | -0.51 | 0.33 |
| Small | 0.37 | 0.19 | 25.1 | -0.01 | 0.75 |

*Table S15. Model estimates (referred to acoustic PC3)*

| behPC1 | Eye peak temperature | SE | df | lower.CL | upper.CL |
|---|---|---|---|---|---|
| 0.39 | -0.19 | 0.13 | 32.3 | -0.44 | 0.07 |
| 1.65 | 0.05 | 0.18 | 24.7 | -0.31 | 0.42 |
| 2.91 | 0.29 | 0.27 | 28.0 | -0.26 | 0.85 |

*Table S16. Model estimates (referred to acoustic PC3)*

| behPC1 | Eye peak temperature | SE | df | lower.CL | upper.CL |
|---|---|---|---|---|---|
| Large | 0.29 | 0.23 | 24.9 | -0.18 | 0.76 |
| Small | -0.18 | 0.20 | 37.1 | -0.58 | 0.22 |

*Table S17. Model estimates (referred to acoustic PC4)*

| behPC1 | Eye peak temperature | SE | df | lower.CL | upper.CL |
|---|---|---|---|---|---|
| 0.39 | -0.04 | 0.10 | 28.1 | -0.24 | 0.17 |
| 1.65 | 0.20 | 0.14 | 20.3 | -0.08 | 0.49 |
| 2.91 | 0.44 | 0.21 | 23.4 | 0.01 | 0.88 |

*Table S18. Post hoc test (referred to acoustic PC1, PC2, PC3 and PC4)*

| | Contrast | estimate | SE | df | t.ratio | p.value |
|---|---|---|---|---|---|---|
| AcPC1 | Large - small | -2.21 | 0.84 | 18.3 | -2.64 | 0.02 |
| AcPC2 | Large - small | -0.46 | 0.23 | 29.8 | -2.00 | 0.05 |
| AcPC3 | Large - small | 0.47 | 0.24 | 45.2 | 1.95 | 0.06 |
| | BehPC1 0.39 - BehPC1 1.65 | -0.24 | 0.11 | 43.9 | -2.14 | 0.09 |

| | | | | | | |
|---|---|---|---|---|---|---|
| | BehPC1 0.39 - BehPC1 2.91 | -0.48 | 0.23 | 43.9 | -2.14 | 0.09 |
| | BehPC1 1.65 - BehPC1 2.91 | -0.24 | 0.11 | 43.9 | -2.14 | 0.09 |
| AcPC4 | BehPC1 0.39 - BehPC1 1.65 | -0.24 | 0.09 | 37.4 | -2.62 | 0.03 |
| | BehPC1 0.39 - BehPC1 2.91 | -0.48 | 0.18 | 37.4 | -2.62 | 0.03 |
| | BehPC1 1.65 - BehPC1 2.91 | -0.24 | 0.09 | 37.4 | -2.62 | 0.03 |
